# Multi-BRCT domain protein Brc1 links Rhp18/Rad18 and γH2A to maintain genome stability during S-phase

**DOI:** 10.1101/150714

**Authors:** Michael C. Reubens, Sophie Rozenzhak, Paul Russell

## Abstract

DNA replication involves the inherent risk of genome instability, as replisomes invariably encounter DNA lesions or other structures that stall or collapse replication forks during S-phase. In the fission yeast *Schizosaccharomyces pombe*, the multi-BRCT domain protein Brc1, which is related to budding yeast Rtt107 and mammalian PTIP, plays an important role in maintaining genome integrity and cell viability when cells experience replication stress. The C-terminal pair of BRCT domains in Brc1 were previously shown to bind phospho-histone H2A (γH2A) formed by Rad3/ATR checkpoint kinase at DNA lesions; however, the putative scaffold interactions involving the N-terminal BRCT domains 1-4 of Brc1 have remained obscure. Here we show that these domains bind Rhp18/Rad18, which is an E3 ubiquitin protein ligase that has crucial functions in postreplication repair. A missense allele in BRCT domain 4 of Brc1 disrupts binding to Rhp18 and causes sensitivity to replication stress. Brc1 binding to Rhp18 and γH2A are required for the Brc1-overexpression suppression of *smc6-74,* which impairs the Smc5/6 structural maintenance of chromosomes complex required for chromosome integrity and repair of collapsed replication forks. From these findings we propose that Brc1 provides scaffolding functions linking γH2A, Rhp18, and Smc5/6 complex at damaged replication forks.

## INTRODUCTION

Genome stability is especially at risk during the DNA synthesis (S)-phase of the cell cycle, when replisomes encounter DNA lesions or chromatin-bound proteins, or they collide with converging replication or transcriptional machinery, potentially leading to replication fork collapse and ensuing deleterious genomic alterations. Faced with the critical requirement for replication fidelity to maintain cell viability and prevent disease (1-3), eukaryotic organisms have evolved a complex and highly regulated network of DNA damage response (DDR) pathways that work in conjunction with the replicative machinery to maintain genome integrity during S-phase (4-8). Thus, DDRs coordinate DNA replication, repair, and cell cycle progression to safeguard the genome.

In *Schizosaccharomyces pombe* (*S. pombe*), as in all eukaryotes, the replication stress response is initiated by the detection of replication protein a (RPA)-coated single-stranded DNA (ssDNA) that forms at stalled or damaged replication forks. This accumulation of RPA-bound ssDNA serves as a signal to activate the master checkpoint kinase Rad3/ATR, which phosphorylates key substrates including an SQ motif on the C-terminal tail of histone H2A in chromatin flanking the stalled or collapsed replication fork (9, 10). Phospho-histone H2A, known as γH2A in yeast and equivalent to γH2AX in mammals, serves as a recruitment platform for key DDR proteins, including Brc1, Crb2, and Mdb1 in *S. pombe* (11-15).

Brc1 is an 878-amino acid (aa) protein that contains four N-terminal and two C-terminal BRCT domains separated by an intervening linker region containing a nuclear localization signal (Figure 1A) (16). This domain organization is shared with *Saccharomyces cerevisiae* Rtt107 and mammalian PTIP, which are important genome protection proteins (17-22). Brc1 was first identified in *S. pombe* as an allele-specific high-copy suppressor of the hypomorphic *smc6-74* allele, which impairs the function of the essential Smc5/6 structural maintenance of chromosomes complex (23). Brc1 is not required for cell viability unless cells are challenged with DNA damaging agents that collapse replication forks, or they are defective in specific processes related to DNA replication. For example, Brc1 is essential for cell viability in mutants lacking Rqh1, which is a RecQ-like DNA helicase involved in DNA replication and repair (9, 24). Rqh1 is homologous to Sgs1 in *S. cerevisiae* and the BLM, WRN and RTS/RECQ4 enzymes in humans that are associated with cancer predisposition and/or premature aging (25). Brc1 is also crucial when loss of deoxycytidylate deaminase (dCMP deaminase) creates imbalanced pools of deoxyribonucleoside triphosphates (dNTPs) required for DNA synthesis and repair (26), or when there are defects in replication factor c (RFC), which loads proliferating cell nuclear antigen (PCNA) clamp onto duplex DNA (10). Perhaps most notably, Brc1 becomes essential in cells with compromised Smc5/6 complex function (23, 24). The reported function of the Smc5/6 complex at collapsed replication forks (27-29), combined with the observed sensitivity of Brc1-deficient cells to agents known to generate lesions and disrupt replication fork progression during S-phase, suggests that Brc1 functions in the response to DNA damage during replication stress (24). Further supporting this idea, Brc1 was shown to localize at sites of replication stress through the interaction of its C-terminal BRCT domains with γH2A (12). Collectively, these data suggest that Brc1 stabilizes stalled replication forks and assists in the repair of collapsed replication forks (9, 10, 12, 16, 30).

**Figure 1:**
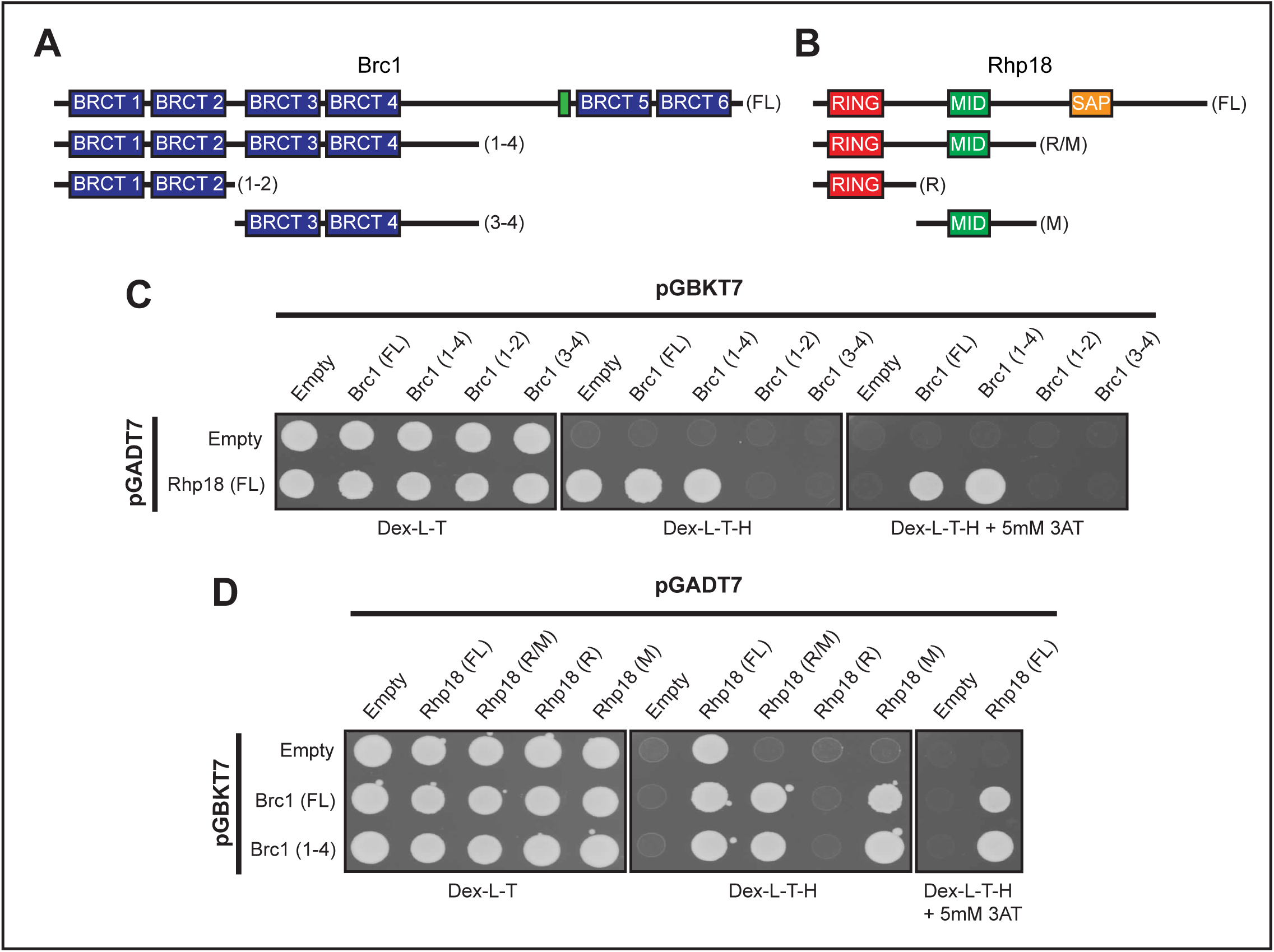
The Brc1-Rhp18 interaction requires the N-terminal BRCT domains of Brc1 and the Mid domain of Rhp18. **A** and **B.** Schematic representations of Brc1 (**A**) and Rhp18 (**B**) fragments tested for physical interactions by yeast two-hybrid analysis (fragments sizes for both Brc1 and Rhp18 are listed in the materials and methods section). The location of Brc1’s nuclear localization signal is depicted by the green box in A. **C.** Yeast two-hybrid results showing full length Brc1 and BRCT domains 1-4 interact with full length Rhp18. Interactions were judged from the Dex-L-T-H+5mM 3AT due to observed one-hybrid activity of full length Rhp18 in pGADT7 on Dex-L-T-H. **D.** Yeast two-hybrid results indicating Rhp18’s Mid domain is sufficient to support the interaction with Brc1. Interactions for full length Rhp18 were evaluated from the Dex-L-T-H+5mM3AT due to observed one-hybrid activity of full length Rhp18 in pGADT7 on Dex-L-T-H. Removal of the SAP domain of Rhp18 alleviated the observed one-hybrid activity, allowing assessment of physical interactions on Dex-L-T-H for the R/M, R, and M fragments of Rhp18.

Exactly how Brc1 protects genomic stability during S-phase has remained elusive. Extensive genetic interaction analysis has established that Brc1 is especially crucial for replication stress resistance and cell viability when DNA replication or other genome protection mechanisms are impaired (9, 10, 23, 24, 31-33); however, lack of specific measureable enzymatic activity and limited protein interaction data have hindered progression in understanding Brc1’s role in the maintenance of genomic stability under these circumstances. Brc1 binds γH2A, but this interaction probably serves to properly localize Brc1 at DNA lesions, where it engages with other proteins that are currently unknown. Moreover, mutations that disrupt Brc1 binding to γH2A only partially impair Brc1 function, indicating γH2A-independent roles for Brc1 that do not absolutely require formation of extensive domains of chromatin-bound Brc1 flanking stalled or collapsed replication forks (12, 16). The *smc6-74* suppression by Brc1 overexpression was shown to require Rhp18, but whether this dependence reflects physical associations between Brc1, Rhp18, or the Smc5/6 holocomplex is unknown (24, 31). Rhp18, known as Rad18 in other organisms, is an E3 ubiquitin protein ligase, which binds RPA-coated ssDNA, where it functions with its cognate E2 enzyme, Rad6, to control the initial steps of post replicative repair (PRR) via proliferating cell nuclear antigen (PCNA) ubiquitination (34).

In this report, we identify Rhp18 as a binding partner for Brc1 and describe a mutation that disrupts this interaction. Binding studies and functional analysis suggest the interaction with Rhp18 is essential for Brc1 overexpression to suppress *smc6-74*. Moreover, Brc1 binding to γH2A is critical when the function of the Smc5/6 complex is impaired. These results suggest that Brc1’s role in promoting genomic stability during S-phase is mediated through coordinated binding with γH2A and Rhp18 at sites of replication stress.

## RESULTS

### Brc1 physically interacts with Rhp18

BRCT domains often mediate physical interactions amongst proteins (35, 36). Brc1 contains six BRCT domains but no documented enzymatic activity, thus it seemed likely that it functions as a scaffold for other DDR proteins at stalled or damaged replication forks. To date the only interacting partner identified for Brc1 has been γH2A (12). Therefore, we sought to identify additional Brc1 interacting proteins through a yeast two-hybrid (Y2H) screen using full length Brc1 as bait. The results from this preliminary screen returned multiple hits for Rhp18, which was notable given that Rhp18 is required for the rescue of *smc6-74* by Brc1 overexpression (24, 31).

To confirm these preliminary results, we cloned full length *brc1* cDNA into *pGBKT7* and full length *rhp18* cDNA into *pGADT7* and assessed their two-hybrid interactions. Rhp18 displayed one-hybrid activity on standard -His selective media (Dex-L-T-H), but addition of 5 mM 3-Amino-1,2,4 Triazole (3-AT) to the media suppressed this activity and confirmed the Brc1-Rhp18 interaction reported from our initial screen (Figure 1C). These results were later validated through co-immunoprecipitation, as described below.

### Identification of domains mediating the Brc1-Rhp18 interaction

With Rhp18 identified as a binding partner of Brc1, we sought to narrow down the protein domains mediating this interaction. Fragments of Brc1 were evaluated for their ability to bind full length Rhp18 using the Y2H method (Figure 1A). This analysis revealed that a Brc1 fragment containing BRCT domains 1-4 plus part of the linker region was sufficient for the Y2H interaction with Rhp18. However, division of this fragment between the BRCT domains 2 and 3 eliminated the Y2H signal, suggesting that BRCT domains 1-4 likely function as a binding module for Rhp18 (Figure 1C).

Rhp18 is a 387-aa protein that contains an N-terminal really interesting new gene (RING) domain, a central Mid zinc finger domain, and a C-terminal SAF-A/B, Acinus, Pias (SAP) domain. The N-terminal RING domain coordinates binding of Rad6, an E2 ubiquitin conjugating enzyme, along with the C-terminal Rad6 binding interface. Aside from coordinating Rad6 binding, the RING domain is known to mediate Rhp18’s E3 ubiquitin ligase activity to initiate PRR pathways (31, 37). The C-terminal SAP domain has been suggested to possess ssDNA-binding activity, which might recruit Rhp18 to sites of DNA lesions in concert with Rhp18’s ability to bind RPA (38-40). However, the function of the central Mid zinc finger domain is uncertain, as some reports have claimed it mediates replication-independent DNA binding (38), while others have shown that this domain functions as a ubiquitin-binding zinc finger (UBZ) domain (37).

To identify the regions of Rhp18 required for its interaction with Brc1, we utilized our Y2H approach. Fragments of Rhp18 (Figure 1B) were cloned into pGADT7 and tested for their ability to interact with full length Brc1 or the Brc1(1-4) fragment expressed from pGBKT7 (Figure 1A). Removal of the SAP domain alleviated the observed one-hybrid activity in pGADT7, and allowed scoring of interactions on restrictive plates lacking 3-AT. The Rhp18(R/M) fragment supported growth on the restrictive plates, suggesting that the SAP domain is not essential for the Brc1 interaction (Figure 1D). The N-terminal RING domain failed to support growth on the restrictive plates when combined with either full length Brc1 or the N-terminal BRCT domains (Figure 1D), suggesting that the RING domain alone is insufficient to mediate the interaction with Brc1. The combined results above implied that the Mid domain alone may be sufficient to interact with Brc1, and the two-hybrid analysis expressing only the Mid domain in pGADT7 in conjunction with the Brc1 fragments in pGBKT7 corroborated that assumption (Figure 1D). Therefore, our results suggest the Mid zinc finger domain of Rhp18 alone is sufficient to support the physical interaction with Brc1.

### Mutation of BRCT domain 4 severely attenuates the Brc1-Rhp18 physical interaction

Having identified the regions of Brc1 and Rhp18 required for their physical interaction, we next turned our attention to analyzing the effects of previously characterized point mutations altering conserved residues in the BRCT domains of Brc1 (12, 16). As expected, the *brc1-T672A* mutation that disrupts binding to γH2A did not diminish binding to Rhp18 (Figure 2B). Brc1 proteins with altered residues in BRCT domain 2 (*G136A* and *TH148, 149SG*) or BRCT domain 3 (*R268K* and *W298F, P301G*) also maintained the Y2H interaction with Rhp18. In contrast, the *brc1-HYP307-9GFG* allele, which alters 3 residues in BRCT domain 4 near the BRCT 3-4 linker, eliminated the Y2H interaction with Rhp18 (Figure 2B). Importantly, previous work showed that *brc1-HYP307-9GFG* impaired Brc1 function in genotoxin resistance without disrupting its ability form γH2A-dependent nuclear foci (16), suggesting that this allele disrupted critical scaffolding properties of Brc1.

**Figure 2.**
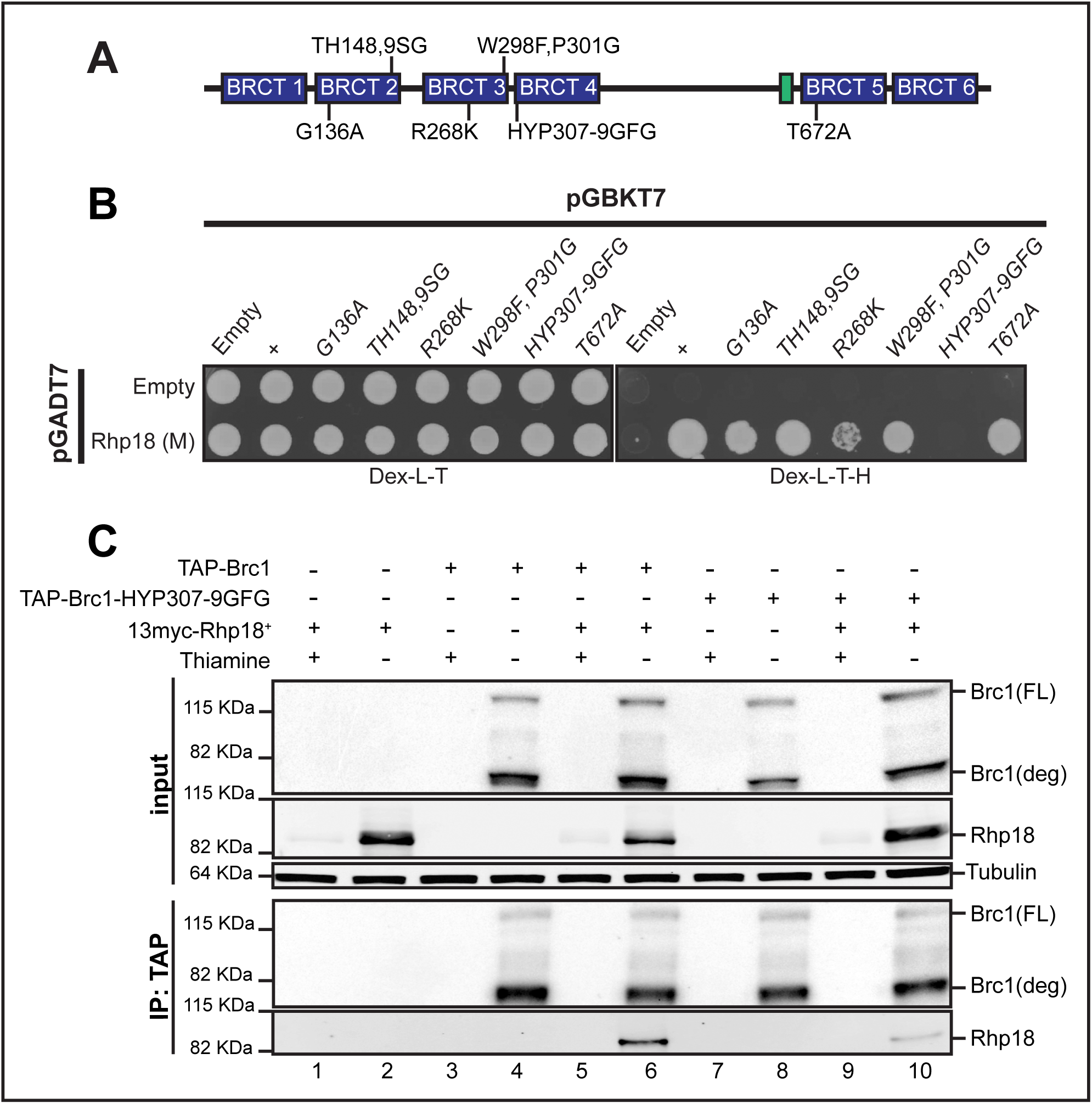
Mutation of BRCT domain 4 attenuates the Brc1-Rhp18 interaction. **A.** A schematic representation of published Brc1 point mutations (Lee and Russell, 2013) tested for physical interaction with the Mid domain of Rhp18 by yeast two-hybrid analysis. **B**. Yeast two-hybrid results indicating that mutation of BRCT4 inhibits the interaction with the Mid domain of Rhp18. **C**. Results from co-immunoprecipitation experiments demonstrating the interaction between full length Brc1 and Rhp18, as well as the reduction in Rhp18 binding observed with TAP-Brc1-HYP307-9GFG versus TAP-Brc1.

To confirm the yeast-two hybrid assays we performed co-immunoprecipitation experiments. Utilizing *nmt41* promoter driven expression of N-terminally tagged Rhp18 and Brc1, we found that 13myc-tagged Rhp18 readily co-precipitated with TAP-tagged Brc1 (Figure 2C, lane 6). In contrast, Rhp18 co-immunoprecipitation with TAP-tagged Brc1*-HYP307-9GFG* was strongly diminished (Figure 2C, lane 10).

### Efficient rescue of *brc1Δ* by expression of *brc1-T672A* but not *brc1-HYP307-9GFG*

With the identification of point mutations of Brc1 that disrupt binding to Rhp18 or γH2A, we directly compared the effects of these mutations in determining cellular resistance to DNA damage. We expressed *brc1^+^*, the Rhp18-binding defective mutant *brc1-HYP307-9GFG*, or the previously described γH2A-binding defective mutant *brc1-T672A* (12), from the moderate strength *nmt41* promoter in plasmid pREP41X. As an additional control, we also expressed *brc1-W298F*, *P301G*, containing mutations in BRCT domain 3, which did not disrupt the Brc1-Rhp18 Y2H interaction (Figure 2B). We found that expression of *brc1^+^*, *brc1-W298F:P301G*, and *brc1-T672A* were all able to fully rescue *brc1Δ* MMS sensitivity (Figure 3A, rows 4, 5, and 7). In contrast, expression of *brc1-HYP307-9GFG* resulted in an extremely weak rescue of the MMS phenotype when compared to the vector only control (Figure 3A, row 6). We obtained essentially the same results when we repeated the experiment in an *htaAQ* genetic background (*hta1-S129A hta2-S128S*) (Figure 3B), which lacks the ability to form γH2A (11). Thus, defects in Brc1 binding to γH2A can be suppressed by Brc1 overexpression; however, the impacts of the Rhp18-binding defective *brc1-HYP307-9GFG* mutation on Brc1 function cannot be compensated for by merely increasing its cellular concentrations.

**Figure 3.**
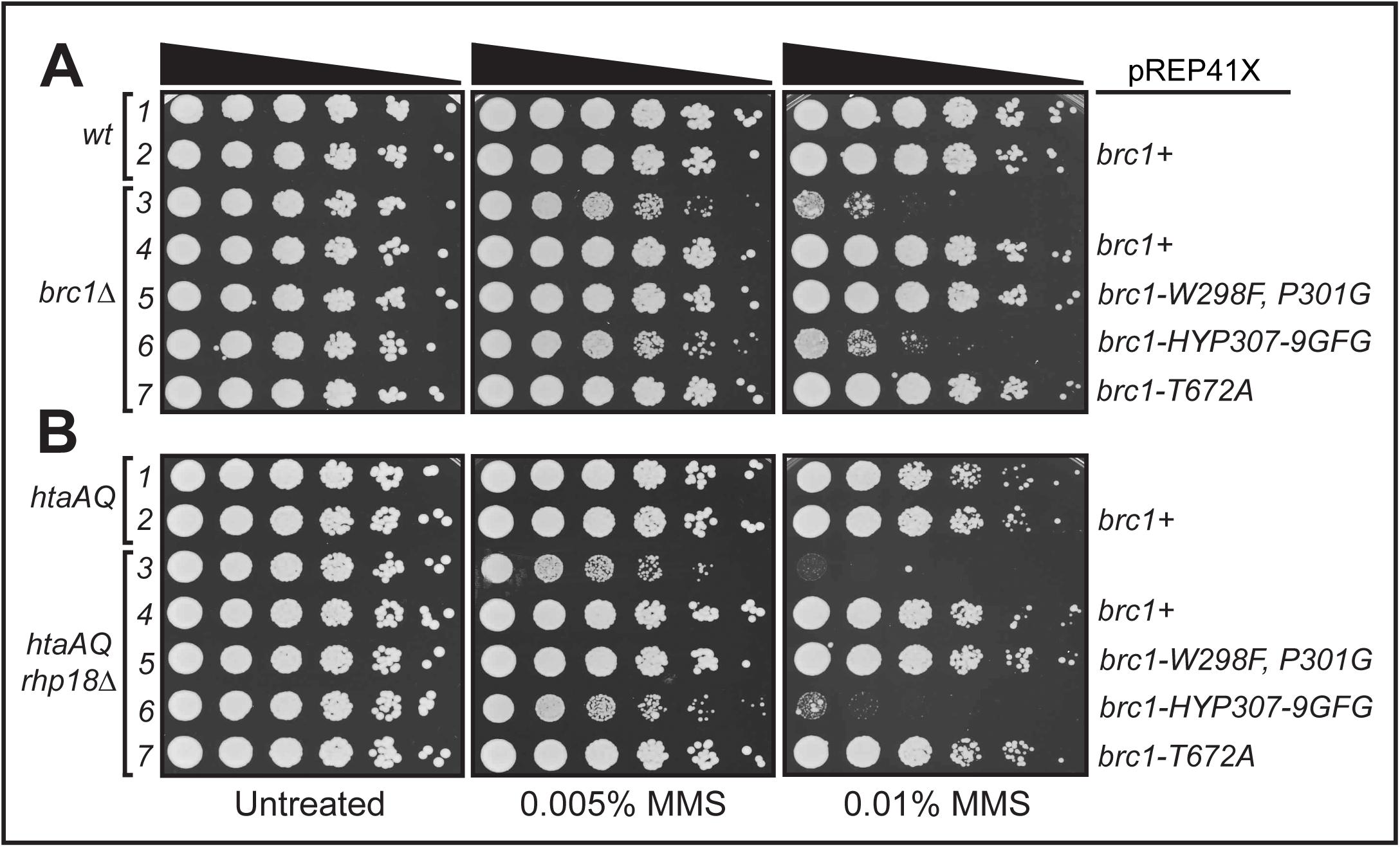
Efficient rescue of brc1Δ by expression of brc1-T672A but not brc1-HYP307-9GFG. **A**. Functional evaluation of four *brc1* alleles in response to MMS treatment in a *brc1Δ* genetic background suggests that *brc1-HYP307-9GFG* (Lee and Russell, 2013) retains more activity than *brc1Δ*, but significantly less than the previously published γH2A binding mutant *brc1-T672A* (Williams, J., *<u>et.al</u>.,* 2010), which rescued *brc1Δ* as well as *brc1^+^* and *brc1-W298F*, *P301G*. **B**. Functional evaluation of the *brc1* alleles in response to MMS treatment in the *htaAQ brc1Δ* genetic background, demonstrating the Brc1-Rhp18 interaction is more essential for Brc1 function in an overexpression situation than its ability to bind γH2A.

### Brc1 binding to Rhp18 and γH2A are important for suppression of *smc6-74* by Brc1 overexpression

We next investigated the relationships between Brc1 binding to γH2A or Rhp18 and its ability to rescue *smc6-74*. Utilizing the same approach as described above, we expressed the *brc1* alleles from *pREP41X* in the *smc6-74* genetic background and tested MMS sensitivity. As seen for the *brc1Δ* rescue experiments, expression of *brc1^+^* or *brc1-W298F, P301G* fully suppressed the *smc6-74* MMS-sensitive phenotype in these assays (Figure 4A, rows 4-5). In contrast, expression of *brc1-HYP307-9GFG* resulted in no suppression of *smc6-74* (Figure 4A, row 6). The correlation of attenuated Rhp18 binding and lack of *smc6-74* suppression observed for *brc1-HYP307-9GFG* overexpression suggests that binding to Rhp18 is critical for Brc1 function. Interestingly, suppression *smc6-74* by *brc1-T672A* overexpression was weakened in comparison to *brc1^+^* overexpression (Figure 4A, row 7), suggesting that γH2A-binding by Brc1 is important for suppression of *smc6-74*.

**Figure 4.**
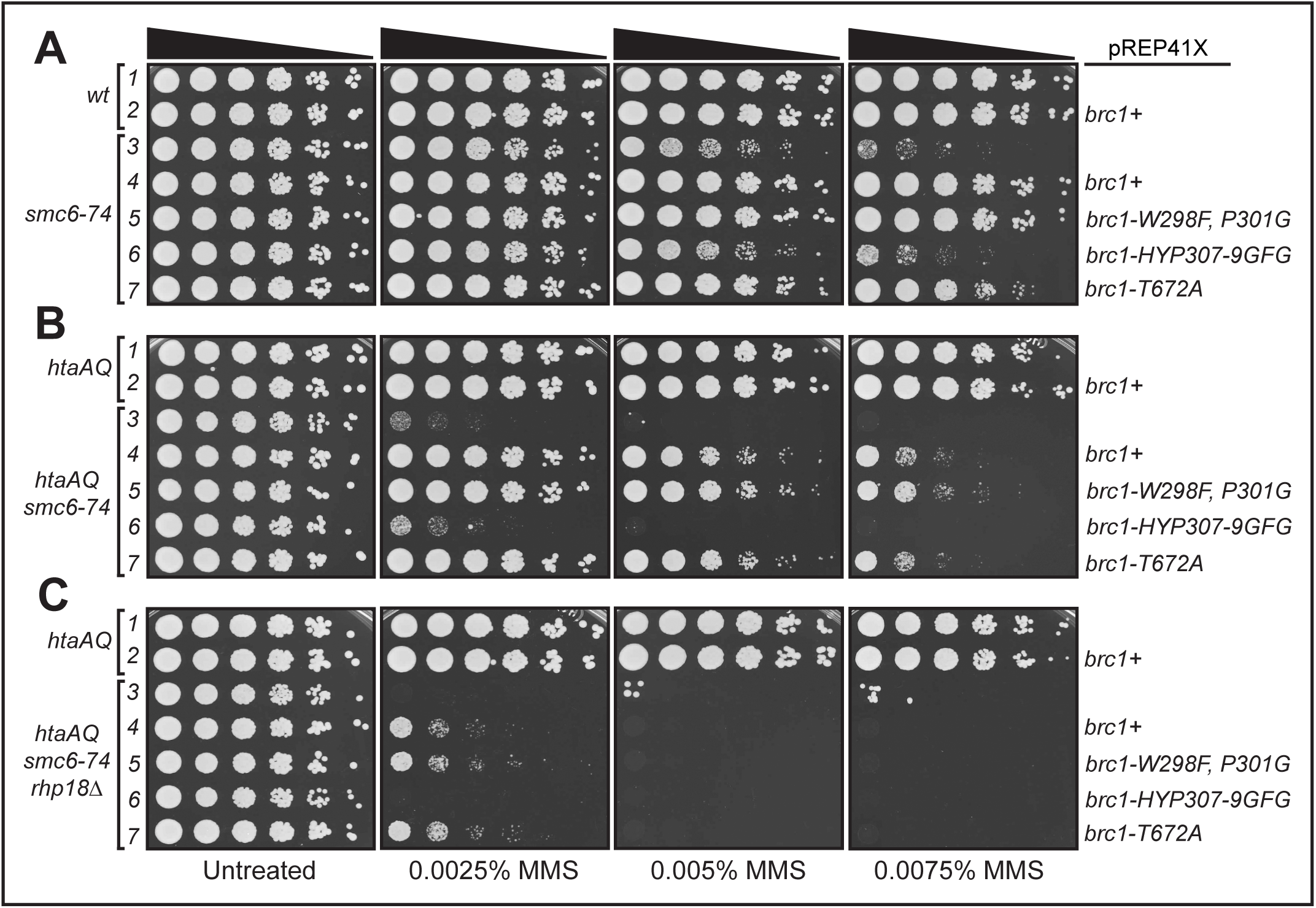
Rhp18 and γH2A binding are required for efficient rescue of *smc6-74* by Brc1 overexpression. **A**. Results from *smc6-74* suppression experiments comparing the rescue efficiency of the four *brc1* alleles. The *brc1-HYP307-9GFG* allele that disrupts Brc1 binding to Rhp18 prevents Brc1 overexpression suppression of *smc6-74*. The *brc1-T672A* mutation that abrogates binding to γH2A impairs Brc1 overexpression suppression of *smc6-74*. **B**. Results from *smc6-74 htaAQ* suppression experiments comparing the four *brc1* mutations, supporting the MMS dose dependence for γH2A binding in mediating the *scm6-74* rescue by Brc1. **C**. Results from *brc1* overexpression assays in the *smc6-74 htaAQ rhp18Δ* background, suggesting the failure of *brc1-HYP307-9GFG* to rescue *smc6-74* is not completely explained by its inability to bind Rhp18.

To further investigate whether binding to γH2A is important for the Brc1 overexpression rescue of *smc6-74*, we assessed the ability or our *brc1* alleles to suppress *smc6-74* in an *htaAQ* background. Importantly, combining the *smc6-74* and *htaAQ* mutations in the same strain caused an apparent synergistic increase in MMS sensitivity (compare Figure 4A, row 3 to Figure 4B, rows 1 and 3). Expression of *brc1^+^* or *brc1-W298F, P301G* suppressed the *smc6-74* MMS-sensitive phenotype, but MMS-resistance was diminished in the *htaAQ* background (compare Figures 4A-B, rows 4-5). As expected, an approximately equal level of suppression was observed for *brc1-T672A* overexpression in the *htaAQ* background (Figure 4B, row 7), and no suppression was observed for *brc1-HYP307-9GFG* overexpression (Figure 4B, row 6).

These results suggest that interactions of Brc1 with γH2A and Rhp18 are important for suppression of *smc6-74* by Brc1 overexpression. To further test this hypothesis, we examined suppression in the *htaAQ rhp18Δ smc6-74* genetic background. As expected, Brc1 overexpression only weakly suppressed MMS sensitivity of *htaAQ rhp18Δ smc6-74* cells (Figure 4C). This very weak suppression effect was observed for *brc1*^+^, *brc1-W298F*, *P301G* and *brc1-T672A* overexpression. However, this weak suppression was eliminated when we over-expressed *brc1-HYP307-9GFG* (Figure 4C, row 6). This result implies that the *smc6-74* suppression defect of *brc1-HYP307-9GFG* is not fully explained by its inability to interact with Rhp18.

## DISCUSSION

Two-thirds of the mutations associated with human cancers arise from DNA replication errors, emphasizing the need to understand how cells protect genome integrity during S-phase (1). Brc1 preserves genomic stability in response to replication stress, but the mechanism has remained elusive (23, 24, 31-33). One well-defined property of Brc1 is its ability to bind γH2A through its C-terminal BRCT domains. In this report, we have identified Rhp18 as another binding partner of Brc1. This interaction is mediated through the N-terminal region of Brc1 containing BRCT domains 1-4 and it was disrupted by the *brc1-HYP307-9GFG* allele containing clustered mutations at the beginning of BRCT domain 4. As observed for wild type Brc1, the Brc1 protein encoded by *brc1-HYP307-9GFG* properly localizes in the nucleus, where it forms foci in response to replication stress (16). Thus, the *brc1-HYP307-9GFG* mutation does not appear to grossly disrupt Brc1 protein stability or localization, or its ability to bind γH2A-marked chromatin flanking stalled or damaged replication forks. From these results, we propose that the *brc1-HYP307-9GFG* mutation most likely disrupts a scaffolding function of Brc1 that involves binding Rhp18. This model is consistent with the requirements for Rhp18 to tolerate genotoxins that cause replication fork stalling and collapse, and the requirement for Rhp18 in suppression of *smc6-74* by Brc1 overexpression (24, 31). Importantly, the *brc1-HYP307-9GFG* mutation abrogates *smc6-74* suppression by Brc1 overexpression.

As a technical note, we used the yeast two-hybrid method for our studies because we could not reliably precipitate full-length Brc1 in non-denaturing buffers. However, as shown in Figure 2C, we have largely solved this problem through the use of an N-terminal TAP tag, although a substantial amount of Brc1 appears to still be proteolytically cleaved. Importantly, we could confirm our two-hybrid findings with these co-immunoprecipitation studies. The ability to precipitate TAP-tagged Brc1 in non-denaturing buffers will make it possible to employ proteomic methods in future experiments, which has been a very profitable strategy for analyzing the function of Rtt107 (21).

We observed that suppression of *smc6-74* MMS sensitivity by Brc1 overexpression is largely ablated when Brc1 cannot bind to γH2A. Moreover, elimination of γH2A strongly sensitizes *smc6-74* cells to MMS. These data strengthen the evidence linking Brc1 to the proposed role for the Smc5/6 complex in homologous recombination (HR)-mediated repair of stalled replication forks (27-29). One possibility is that diminished Smc5/6 function creates a greater demand for Brc1 to act in fork stabilization and potential channeling of fork repair, or resolution, through alternate pathways that likely depend on Mus81-Eme1 or Slx1-Slx4 (24, 41, 42).

It has been reported that Rhp18 is recruited to ssDNA and RPA-bound ssDNA (39, 40). Given that RPA bound to ssDNA is sensed by Rad3/ATR, which phosphorylates H2A to form γH2A (5, 43), it is unlikely that Rhp18 localization at DNA lesions requires binding to Brc1. However, the potential presence of Rhp18 at the site of stalled replication forks, through its interaction with RPA, could provide a potential binding surface at the fork to explain the γH2A-independent function for Brc1 that has been suggested in previous publications (12, 16).

It was previously reported that the dependence of the *smc6-74* rescue on Rhp18 was due to a requirement for the translesion synthesis (TLS) branch of PRR at higher MMS doses; however, the requirement at lower MMS concentrations could not be attributed to a known function of Rhp18 (24, 31). The results presented here combined with previously published data suggesting that RPA-coated ssDNA can negatively regulate Rad51 strand invasion (44-46), suggest the potential for the binding of Brc1 to RPA-bound Rhp18 to stabilize RPA on its ssDNA substrate, thus potentially inhibiting HR-mediated fork resolution. In the presence of a low level of DNA alkylation damage this stabilization of RPA could serve to inhibit Rad51 strand invasion, therefore inhibiting the onset of HR-mediated fork resolution and allowing excision pathways to repair alkylated bases. This idea would be consistent with the requirement for Rhp18 in the *smc6-74* rescue in response to low level MMS treatment, accompanied by no known Rhp18 activity under those circumstances (31). Furthermore, the interaction between Brc1 and RPA bound Rhp18 would allow for the process of TLS to be mediated at MMS doses that can be tolerated by this method of PRR. Under this scenario, it is also possible that in response to high accumulation of alkylated bases that cannot be repaired via the previously mentioned mechanisms, the fork could be held in a stable confirmation while Brc1 mediates the firing of surrounding dormant replication origins in an Orc1 dependent manner (47), allowing the resolution of the stalled fork via HR-mediated pathways late in S-phase or possibly in G2. If this hypothetical series of events are correct it could potentially explain the complex epistatic relationships between Brc1 and factors involved in multiple DDR pathways (23, 24, 31-33).

Finally, we note that a recent proteomics study with human cells led to the discovery of a putative DNA repair factor, consisting of SLF1 and SLF2, which physically links Smc5/6 complex to Rad18 bound to RNF168-catalyzed ubiquitin chains at certain types of DNA lesions (48). SLF1, also known as BRCTx, is a multi-BRCT domain protein that uses its BRCT domains to bind Rad18 (49, 50). Indeed, this SLF1/BRCTx-Rad18 interaction was first discovered through a two-hybrid screen that used BRCTx as bait, just as we discovered the Brc1-Rhp18 interaction in fission yeast. The protein interaction network involving SLF1/SLF2, Rad18 and Smc5/6 complex does not include Rad6, suggesting that Rad18 functions structurally and not catalytically in this network (48). Strikingly, suppression of *smc6-74* by Brc1 overexpression on low dose MMS does not require Rhp6, the fission yeast ortholog of Rad6 (31). Rtt107, the Brc1-like protein in budding yeast, was shown to mediate Smc5/6 recruitment to DSBs (51). Thus, there appear to be several striking parallels of protein interactions involving Brc1/Rtt107/PTIP BRCT domain proteins, Rad18/Rhp18, and the Smc5/6 complex. We look forward to learning whether these similarities reflect evolutionary divergence of a conserved mechanism of localizing Smc5/6 complex to DNA lesions.

## MATERIALS AND METHODS

### *S. pombe* cultivation and general methods

Standard *S. pombe* methods were conducted as previously described (52), and all *S. pombe* strains used in this study are listed in Table 1. The *rhp18Δ* strain was generated using a targeting construct that replaced the entire *rhp18*^+^ open reading frame with a hygromycin B (*hphMX6*) cassette, and Rhp18 was N-terminally tagged at its endogenous locus using pFA6-*natMX6*-*p41nmt*-*13myc* (described below) using described methods (53). Both the deletion of Rhp18 and the epitope tag were subsequently verified by PCR and sequencing before use in any experiments. Double and triple mutants were generated using random spore analysis, the resulting genotypes were validated by growth on appropriate selective media, and then subsequently verified by PCR.

**Table 1:** Table of strains generated for and used in this study

| Lab ID | Genotype (all strains are leu1-32 and ura4-D18) |
| --- | --- |
| MR5456 | *h<sup>-</sup> + pREP41X* |
| MR5457 | *h<sup>-</sup> + pREP41Xbrc1<sup>+</sup>* |
| MR5458 | *h<sup>-</sup> brc1::kanMX6 + pREP41X* |
| MR5459 | *h<sup>-</sup> brc1::kanMX6 + pREP41Xbrc1<sup>+</sup>* |
| MR5460 | *h<sup>-</sup> brc1::kanMX6 + pREP41Xbrc1-W298F, P01G* |
| MR5461 | *h<sup>-</sup> brc1::kanMX6 + pREP41Xbrc1-HYP307-9GFG* |
| MR5462 | *h<sup>-</sup> brc1::kanMX6 + pREP41Xbrc1-T672A* |
| MR5464 | *h<sup>-</sup> smc6-74 + pREP41X* |
| MR5465 | *h<sup>-</sup> smc6-74 + pREP41Xbrc1<sup>+</sup>* |
| MR5466 | *h<sup>-</sup> smc6-74 + pREP41Xbrc1-W298F, P01G* |
| MR5467 | *h<sup>-</sup> smc6-74 + pREP41Xbrc1-HYP307-9GFG* |
| MR5468 | *h<sup>-</sup> smc6-74 + pREP41Xbrc1-T672A* |
| MR5469 | *h<sup>-</sup> hta1-S129A:ura4<sup>+</sup> hta2-S128A:his3<sup>+</sup> his3-D1 + pREP41X* |
| MR5470 | *h<sup>-</sup> hta1-S129A:ura4<sup>+</sup> hta2-S128A:his3<sup>+</sup> his3-D1 + pREP41XBrc1<sup>+</sup>* |
| MR5471 | *h<sup>-</sup> hta1-S129A:ura4<sup>+</sup> hta2-S128A:his3<sup>+</sup> brc1::kanMX6 his3-D1 + pREP41X* |
| MR5472 | *h<sup>-</sup> hta1-S129A:ura4<sup>+</sup> hta2-S128A:his3<sup>+</sup> brc1::kanMX6 his3-D1 + pREP41XBrc1<sup>+</sup>* |
| MR5473 | *h<sup>-</sup> hta1-S129A:ura4<sup>+</sup> hta2-S128A:his3<sup>+</sup> brc1::kanMX6 his3-D1 + pREP41XBrc1-W298F, P301G* |
| MR5474 | *h<sup>-</sup> hta1-S129A:ura4<sup>+</sup> hta2-S128A:his3<sup>+</sup> brc1::kanMX6 his3-D1 + pREP41XBrc1-HYP307-9GFG* |
| MR5475 | *h<sup>-</sup> hta1-S129A:ura4<sup>+</sup> hta2-S128A:his3<sup>+</sup> brc1::kanMX6 his3-D1 + pREP41XBrc1-T672A* |
| MR5477 | *h<sup>-</sup> hta1-S129A:ura4<sup>+</sup> hta2-S128A:his3<sup>+</sup> smc6-74 his3-D1 + pREP41X* |
| MR5478 | *h<sup>-</sup> hta1-S129A:ura4<sup>+</sup> hta2-S128A:his3<sup>+</sup> smc6-74 his3-D1 + pREP41XBrc1<sup>+</sup>* |
| MR5479 | *h<sup>-</sup> hta1-S129A:ura4<sup>+</sup> hta2-S128A:his3<sup>+</sup> smc6-74 his3-D1 + pREP41XBrc1-W298F, P301G* |
| MR5480 | *h<sup>-</sup> hta1-S129A:ura4<sup>+</sup> hta2-S128A:his3<sup>+</sup> smc6-74 his3-D1 + pREP41XBrc1-HYP307-9GFG* |
| MR5481 | *h<sup>-</sup> hta1-S129A:ura4<sup>+</sup> hta2-S128A:his3<sup>+</sup> smc6-74 his3-D1 + pREP41XBrc1-T672A* |
| MR5485 | *h<sup>-</sup> hta1-S129A:ura4<sup>+</sup> hta2-S128A:his3<sup>+</sup> smc6-74 rhp18::hphMX6 his3-D1 + pREP41X* |
| MR5486 | *h<sup>-</sup> hta1-S129A:ura4<sup>+</sup> hta2-S128A:his3<sup>+</sup> smc6-74 rhp18::hphMX6 his3-D1 + pREP41XBrc1<sup>+</sup>* |
| MR5487 | *h<sup>-</sup> hta1-S129A:ura4<sup>+</sup> hta2-S128A:his3<sup>+</sup> smc6-74 rhp18::hphMX6 his3-D1 + pREP41XBrc1-W298F, P301G* |
| MR5488 | *h<sup>-</sup> hta1-S129A:ura4<sup>+</sup> hta2-S128A:his3<sup>+</sup> smc6-74 rhp18::hphMX6 his3-D1 + pREP41XBrc1-HYP307-9GFG* |
| MR5489 | *h<sup>-</sup> hta1-S129A:ura4<sup>+</sup> hta2-S128A:his3<sup>+</sup> smc6-74 rhp18::hphMX6 his3-D1 + pREP41XBrc1-T672A* |
| MR5490 | *h<sup>-</sup> natMX6:p41-13myc:Rhp18 brc1::kanMX6* |
| MR5491 | *h<sup>-</sup> brc1::kanMX6 + pREP41-N-TAPbrc1<sup>+</sup>* |
| MR5492 | $h^-$ *natMX6:p41-13myc:Rhp18 brc1::kanMX6 + pREP41-N-TAPBrc1<sup>+</sup>* |
| MR5493 | $h^-$ *brc1::kanMX6 + pREP41-N-TAPbrc1-HYP307-9GFG* |
| MR5494 | $h^-$ *natMX6:p41-13myc:Rhp18 brc1::kanMX6 + pREP41-N-TAPBrc1-HYP307-9GFG* |

For immunoprecipitation experiments, exponentially growing cultures of indicated strains were cultivated in appropriately supplemented Edinburgh Minimal Media (EMM2) in the presence, or absence, of 5 μg/mL thiamine for 25 hours at 30°C to actively regulate the expression from the *nmt41* promoter. For MMS survival assays the indicated strains were cultivated in appropriately supplemented EMM2 media in the presence of 5 μg/mL thiamine for 25 hours, log phase cultures were then suspended to 0.4 OD_600_ and serially diluted five-fold onto yeast extract, glucose, and supplements (YES) agar plates containing the designated concentration of MMS. Cell growth was evaluated after 4 days at 30°C as previously described (24, 31).

### Plasmid construction

For the Y2H analysis constructs, all *brc1* fragments (**FL**: 1-878aa, **1-4**: 1-553aa, **1-2**: 1-202aa, and **3-4**: 195-553aa), and point mutants, were isolated using standard PCR methods with NdeI linkers on upstream primers and BamHI linkers on downstream primers. All *rhp18* fragments (**FL**: 1-387aa, **R/M**: 1-202aa, **R**: 1-117aa, and **M**: 111-202aa) were generated using a similar PCR mediated strategy except for using XmaI linkers on the downstream primers. The resulting *brc1* inserts were then ligated into NdeI and BamHI digested pGBKT7, and the *rhp18* fragments were ligated into NdeI and XmaI digested pGADT7. Expression of TAP-Brc1 was achieved by cloning full length *brc1^+^*, or *brc1*-*HYP307-9GFG*, cDNA into NdeI and BamHI digested pREP41-NTAP as previously described (54). For MMS survival assays pREP41X*brc1^+^* was used to rescue the MMS phenotypes as previously described (24, 31), and all evaluated *brc1* point mutants were generated by site-directed mutagenesis (Agilent Technologies) using pREP41X*brc1^+^* as template. To N-terminally tag Rhp18 at its endogenous locus pFA6a-natMX6-p41*nmt*-13myc was constructed by cleaving the 3XFLAG from pFA6a-*natMX6-p41nmt-3XFLAG* (55) and replacing it with the 13myc tag, without its stop codon, isolated from pFA6a-*13myc-natMX6* (53). All plasmids generated for use in this study were sequence verified before use.

### Yeast two-hybrid analysis

All Brc1 and Rhp18 fusion constructs were generated as described above. The resulting fusion protein constructs were transformed into the *S. cerevisiae* AH109 reporter strain (Clontech Matchmaker^®^ Gold System), and co-transformants were selected for by plating on Dex-L-T media. Y2H analysis was carried out by diluting the indicated log phase cultures to an OD_600_ of 0.4 and then spotting them onto restrictive and permissive plates. Control growth was evaluated on Dex-L-T and protein interactions were scored either Dex-L-T-H with, or without, 3-AT based on the presence of one-hybrid activity. All Y2H growth was scored after three days of growth at 32°C.

### Immunoblotting and immunoprecipitation

Whole cell extracts (WCE) were generated from 15 mL cultures of the indicated strains cultivated as described above. Cell pellets were lysed in lysis buffer (50 mM Tris pH 8.0, 150mM NaCl, 5 mM EDTA, 10% glycerol, 0.1% Nonidet-P40, 1 mM NaF, 1 mM PMSF, and Complete Protease Inhibitors) using a FastPrep^®^-24 (MP Biomedicals) following the manufacturer’s protocol. For each lysate 1.5 mg total protein was incubated with rabbit IgG (Sigma) conjugated Dynabeads^®^ M-280 Tosylactivated (Thermo Fisher Scientific) for three hours at 4°C with rotation. The beads were collected and washed three times in lysis buffer before eluting the proteins from the beads by boiling in 1xSDS-PAGE loading buffer (100mM Tris pH6.8, 4% SDS, 20% glycerol, and 0.2% Bromophenol blue). Proteins were resolved on Novex WedgeWell 4-20% Tris-Glycine gels (Thermo Fisher Scientific), transferred via iBlot2^®^ (Thermo Fisher Scientific) to nitrocellulose membranes, and blocked and probed using standard techniques and manufacturer’s recommended protocols. TAP-Brc1 was detected using Peroxidase anti-peroxidase (PAP) soluble complex antibody produced in rabbit (P12291, Sigma-Aldrich) diluted 1:2,000, 13myc-Rhp18 was detected using anti-myc antibody (9E10, Covance) diluted 1;1,000, and tubulin was detected using monoclonal anti-α-Tubulin antibody produced in mouse (T5168, Sigma-Aldrich) diluted 1:10,000.

## ACKNOWLEDGEMENTS

We thank Matthew O’Connell for plasmids and technical insights, Oliver Limbo for technical support, and members of the Russell laboratory and Nick Boddy for helpful discussions.

This research was supported by NIH grants GM059347, CA077325 and CA117638 to P.R.

We declare no conflicts of interest.

